# Altered brain function in anorexia nervosa and bulimia nervosa: A hierarchical series of task-based fMRI meta-analyses

**DOI:** 10.1101/550301

**Authors:** Bangshan Liu, Jin Liu, Yumeng Ju, Mi Wang, Tiebang Liu, Yan Zhang, Lingjiang Li, Marc N. Potenza, Daniel S. Barron

## Abstract

**Background:** Many structural and functional magnetic resonance imaging (MRI) studies have reported differences in brain functional anatomy associated with eating disorders (EDs). We aimed to quantitatively synthesize the current literature of anorexia nervosa (AN) and bulimia nervosa (BN) with a goal of deriving a consensus across these studies.

**Methods:** We performed a hierarchical series of 49 activation likelihood estimation meta-analyses of 101 experiments from 63 studies at the *disorder (AN+BN)*, *diagnosis (AN or BN)*, and *task* (food-, body-, emotion- or cognitive function-related tasks) levels. We further performed sub-analyses at the diagnosis level to assess the influence of disease stage (current or recovered AN), subtype (restrictive AN, rAN), psychiatric comorbidity, medication, and data processing.

**Results:** We did not observe consistent differences in brain activity across all hierarchies. Rather, we observed differences in brain activity in the right fusiform, left inferior parietal lobule and left precuneus that were primarily related to food (representing 19 experiments) and emotion (25 experiments) tasks performed by current AN patients (56 experiments).

**Conclusions:** The task-based functional MRI literature in AN and BN represents a heterogenous set of tasks and patients. Given this heterogeneity, we found very limited convergence across a rather large literature. Such limited convergence suggests individual task-based studies of EDs should be interpreted cautiously. We recommend that future studies of EDs carefully characterize patients based on nutritional status and that, beyond clinical diagnosis, studies utilize a trait- or cognitive-domain-based approach to define their populations of interest.

**Meta-ananlysis Registration:** “Meta-analysis of functional magnetic resonance imaging studies of eating disorders”, https://www.crd.york.ac.uk/prospero (Registration Number: CRD42018086497).

## Introduction

Anorexia nervosa (AN) and bulimia nervosa (BN) are two major diagnostic categories of eating disorders (EDs) characterized by abnormal eating behavior and distorted body image perception (1). The mortality rates associated with EDs are among the highest among all psychiatric disorders, with at least 1 person dying as a direct result of an ED every 62 minutes (2). However, despite their high disease burden, the pathophysiology of AN/BN remains unclear.

Structural and functional magnetic resonance imaging (MRI) studies have reported differences in the brain’s functional anatomy associated with EDs. Some structural MRI studies have shown differences in overall frontal and subcortical volumes as well as in regional cingulate and cerebellar structure (3). However, these results may reflect nutritional and hydration status as structural differences often normalize as patients with EDs recover from malnourishment. What may be important for staging EDs, therefore, is placing patients on a continuum of nourishment and functional capacity instead of focusing solely on macrostructural differences.

Functional MRI studies have reported ED-related differences in brain activity in limbic and fronto-striato-thalamic areas/circuits (e.g., in prefrontal cortex (PFC), thalamus, amygdala, insula, and hippocampus), inferior parietal lobule (IPL) and precuneus. These differences were observed during processing of food-(4), body-(5) or emotion-/social-related (6) stimuli or while performing neurocognitive tasks (7, 8). To derive a quantitative consensus among these studies, Zhu and colleagues (9) conducted a meta-analysis of 21 task-based functional magnetic imaging (fMRI) experiments, many of which assessed the cognitive processing of food- or body-related stimuli. They found that AN was associated with hyperactivation in emotion-related regions (frontal, temporal, insula, caudate, uncus) and hypoactivation in parietal regions, relative to healthy comparison subjects (HCs), when processing food- and body-related stimuli.

The fMRI literature has grown considerably since Zhu et al’s (2012) meta-analysis. Tasks employing emotional and social stimuli or assessing neurocognitive function have become more popular. These additional data give the field the opportunity to again look for consensus across the fMRI literature, this time in a more refined and task-specific manner. Surprisingly, no meta-analysis specific to BN has been published.

Meta-analytic methods continue to evolve in response to the neuroimaging community’s needs. For example, meta-analytic methods have been created to control for possible effects of multiple neighboring foci within one experiment (10) and cluster-level family-wise error (FWE) rate has been adopted by many as the standard correction for multiple comparisons (11). These new meta-analytic considerations are directly relevant to our analysis of the now-expanded data in EDs.

Here we present a comprehensive meta-analysis of 101 task-based fMRI experiments of AN and BN. The primary goal of our study was to quantitatively synthesize the current literature in AN/BN. To accomplish this, we performed 26 meta-analyses organized in a hierarchical manner: the lowest hierarchy (large-scale analyses) combined all studies of ED (AN+BN, *disorder-*level); the middle hierarchy separated studies of AN from BN (*diagnosis-*level); and the highest hierarchy further separated diagnosis-level studies into groups of specific tasks (*task-*level, see Figure 1 and Table 1 for breakdown). A secondary goal of our study was to better understand how disease stage, ED subtypes, psychiatric comorbidity, and medication may influence brain activations. To accomplish this, at the diagnosis-level, we performed 23 additional meta-analyses that grouped population-specific factors (current or recovered ED, ED subtypes, psychiatric comorbidity, and medication status; see Table 2 for breakdown). Therefore, in total we performed 49 meta-analyses to understand more fully the neural correlates of EDs.

**Table 1.**
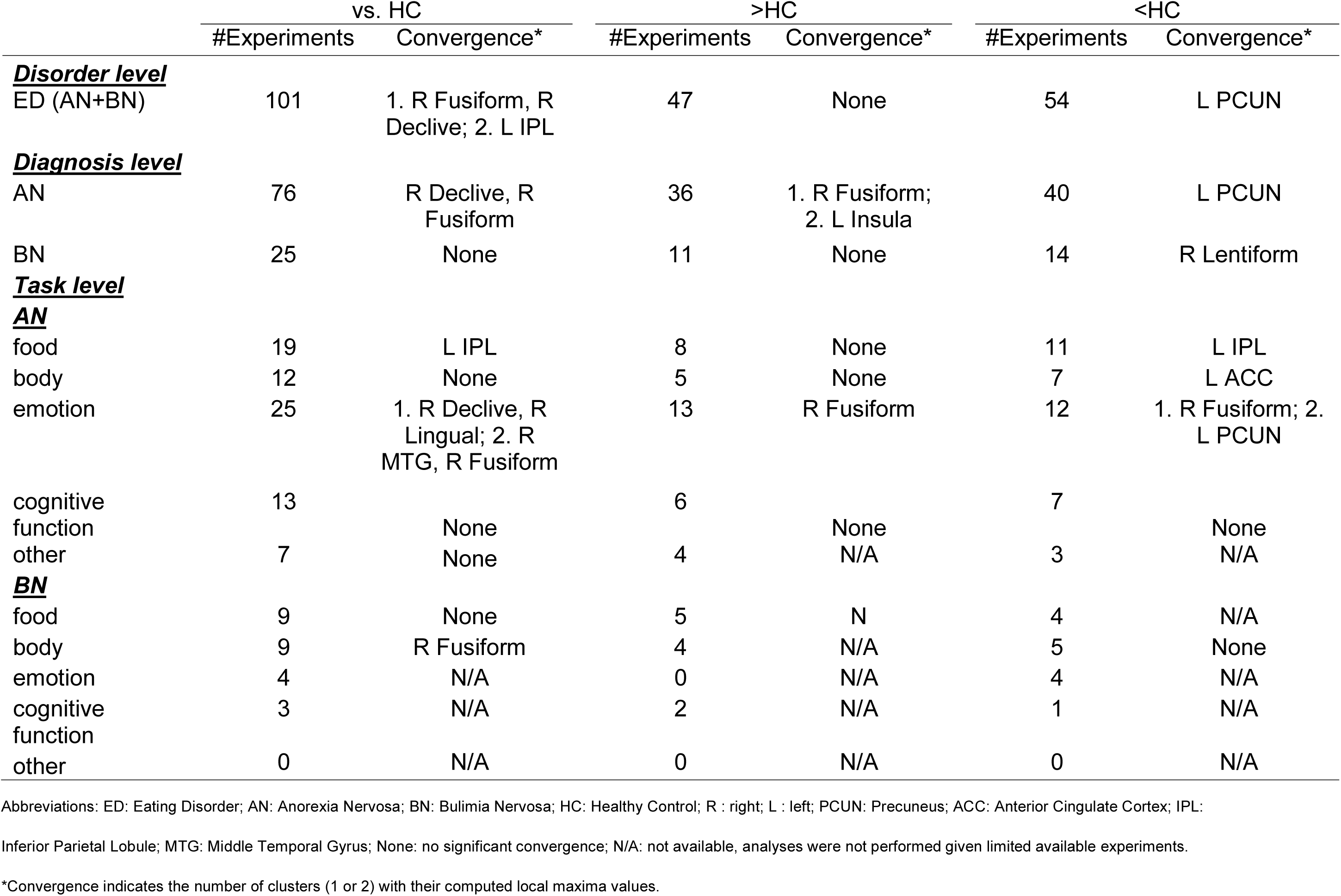
Summary of 26 hierarchical meta-analyses. Results are presented as the Talairaich Daemon anatomical label for the (x,y,z) center of mass for each Activation Likelihood Estimation (ALE) cluster. To condense these results, xyz location, maximum ALE scores and ALE cluster volume is presented in Table S3 of the Supplementary Materials. Abbreviations: ED: Eating Disorder; AN: Anorexia Nervosa; BN: Bulimia Nervosa; HC: Healthy Control; R: right; L: left; PCUN: Precuneus; ACC: Anterior Cingulate Cortex; IPL: Inferior Parietal Lobule; MTG: Middle Temporal Gyrus; N: no significant finding; N/A: not available. *Convergence indicates the number of clusters (1 or 2) with their computed local maxima values.

|  | vs. HC |  | >HC |  | <HC |  |
| --- | --- | --- | --- | --- | --- | --- |
|  | #Experiments | Convergence* | #Experiments | Convergence* | #Experiments | Convergence* |
| **<u>Disorder level</u>** |  |  |  |  |  |  |
| ED (AN+BN) | 101 | 1. R Fusiform, R Declive; 2. L IPL | 47 | None | 54 | L PCUN |
| **<u>Diagnosis level</u>** |  |  |  |  |  |  |
| AN | 76 | R Declive, R Fusiform | 36 | 1. R Fusiform; 2. L Insula | 40 | L PCUN |
| BN | 25 | None | 11 | None | 14 | R Lentiform |
| **<u>Task level</u>** |  |  |  |  |  |  |
| **<u>AN</u>** |  |  |  |  |  |  |
| food | 19 | L IPL | 8 | None | 11 | L IPL |
| body | 12 | None | 5 | None | 7 | L ACC |
| emotion | 25 | 1. R Declive, R Lingual; 2. R MTG, R Fusiform | 13 | R Fusiform | 12 | 1. R Fusiform; 2. L PCUN |
| cognitive function | 13 | None | 6 | None | 7 | None |
| other | 7 | None | 4 | N/A | 3 | N/A |
| **<u>BN</u>** |  |  |  |  |  |  |
| food | 9 | None | 5 | N | 4 | N/A |
| body | 9 | R Fusiform | 4 | N/A | 5 | None |
| emotion | 4 | N/A | 0 | N/A | 4 | N/A |
| cognitive function | 3 | N/A | 2 | N/A | 1 | N/A |
| other | 0 | N/A | 0 | N/A | 0 | N/A |

Inferior Parietal Lobule; MTG: Middle Temporal Gyrus; None: no significant convergence; N/A: not available, analyses were not performed given limited available experiments.

**Table 2.** Summary of 23 diagnosis-level sub-meta-analyses. Results are presented as the Talairaich Daemon anatomical label for the (x,y,z) center of mass for each Activation Likelihood Estimation (ALE) cluster. Coordinate location (as xyz), maximum ALE scores and ALE cluster volume is presented in Table S4 of the Supplementary Materials. Abbreviations: AN: Anorexia Nervosa; BN: Bulimia Nervosa; HC: Healthy Control; rAN: restrictive AN; R: right; L: left; SFG: Superior Frontal Gyrus; MFG: Medial Frontal Gyrus; PCUN: Precuneus; ACC: Anterior Cingulate Cortex; IOG: Inferior Occipital Gyrus. N: no significant finding; N/A: not available. *Convergence indicates the number of clusters (1 or 2) with their computed local maxima values.

|  | vs. HC |  | >HC |  | <HC |  |
| --- | --- | --- | --- | --- | --- | --- |
|  | #Experiments | Convergence* | #Experiments | Convergence* | #Experiments | Convergence* |
| Current AN | 56 | R Declive, R Lingual | 26 | 1. R Fusiform; 2. L Insula | 30 | None |
| Recovered AN | 20 | None | 10 | None | 10 | None |
| rAN | 31 | None | 16 | None | 15 | R SFG |
| Comorbidity-free AN | 29 | R MFG | 14 | R MFG | 15 | L PCUN |
| Medication-free AN | 17 | R Lentiform Nucleus (Putamen) | 9 | None | 8 | 1. R/L ACC, L MFG; 2. L Cingulate gyrus |
| Corrected AN studies | 61 | R Declive, R Fusiform | 29 | 1. R Fusiform, R IOG; 2. R MFG | 32 | L PCUN |
| Comorbidity-free BN | 8 | R Caudate | 4 | N/A | 4 | N/A |
| Medication-free BN | 6 | None | 2 | N/A | 4 | N/A |
| Corrected BN studies | 19 | None | 8 | None | 11 | None |

PCUN: Precuneus; ACC: Anterior Cingulate Cortex; IOG: Inferior Occipital Gyrus. None: no significant convergence; N/A: not available, analyses were not performed given limited
available experiments.. \*Convergence indicates the number of clusters (1 or 2) with their computed local maxima values.

**Figure 1.**
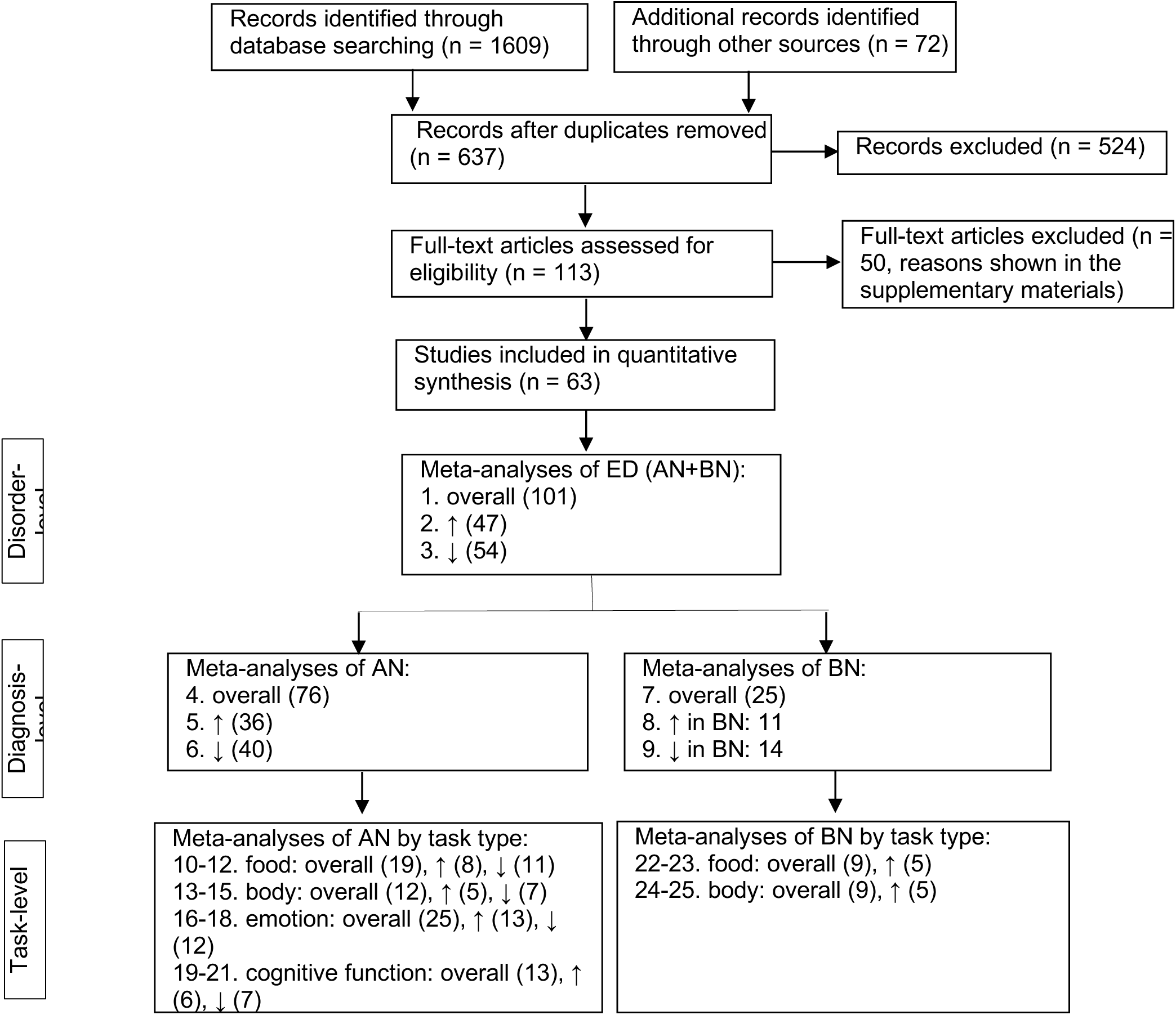
Flowchart of literature screening and hierarchical ALE meta-analyses. Overall, ↑ and ↓indicate vs. HC, >HC and <HC, respectively. The numbers in the leading part of each line are serial numbers of meta-analyses. The numbers in the parentheses are the number of experiments in each meta-analyses. The diagnosis-level sub-analyses are not shown in this figure, which may be referenced in Table 2 and Figure 3.

## Methods and Materials

### Literature search strategy

We searched the Pubmed, Embase, Web of Science, PsycINFO, China National Knowledge Infrastructure (CNKI), Chinese Biomedical Literature (CBM) and WANFANG databases using the terms *eating disorders, anorexia nervosa, bulimia nervosa and magnetic resonance imaging.* A detailed description of the search strategy is described in the supplementary materials. This study was prospectively registered at https://www.crd.york.ac.uk/prospero (CRD42018086497).

### Inclusion and exclusion criteria

We included studies that investigated AN and/or BN with HCs in task-related fMRI paradigms, and that reported significant differences between groups in brain activity in a standard stereotactic coordinate space (Talairach & Torneaux or Montreal Neurological Institute (MNI)).

We excluded studies that:

1) used neuroimaging techniques other than task-based fMRI;

2) restricted the analysis to particular regions, such as region-of-interest (ROI) or seed-based analyses or other mask-based techniques;

3) had a sample size less than 5 in any group;

4) had no HC group or did not report x-y-z foci with effect sizes.

### Data extraction

Of the included studies, we extracted the information about journal, publication year, demographic information (age, sex, adult/adolescent), AN or BN subtypes, current psychiatric comorbidity, current medication status, task type, stimuli used, correction for multiple comparisons, contrasts and coordinates from the papers or through contacting the corresponding authors. A summary of the author-contacting information is shown in Table S1 in the Supplementary Materials. We transformed all x-y-z coordinates to MNI-152 template space prior to analysis.

### Analytic plan

In total, we performed 49 meta-analyses: 26 meta-analyses organized into three hierarchies at the disorder, diagnosis, and task levels; and 23 sub-meta-analyses specific to the diagnosis level. First, we conducted *disorder-level* meta-analyses that pooled AN and BN studies across all tasks, aiming to examine consistent brain activity alteration, hyperactivation or hypoactivation in both EDs across different tasks.

Second, we performed meta-analyses restricted to experiments with only AN or BN participants across all tasks to detect *diagnosis-level* convergence of brain activity alteration, hyperactivation or hypoactivation. We performed sub-meta-analyses to assess possible influences of clinical variables, including disease stage (current (AN-C) or recovered (AN-R)), ED subtypes (restrictive AN (rAN)), current psychiatric comorbidity (comorbidity-free AN/BN), current medication status (medication-free AN/BN) and fMRI methodology (correction for multiple comparisons (corrected AN/BN)) on the findings of diagnosis-level meta-analyses through *sub-analyses* of specific experiments.

Third, we conducted *task-level* meta-analyses to reveal the convergence of functional brain alterations implicated in specific tasks. We categorized 5 types of tasks based on their stimuli and/or designs: related to food, body, emotion, cognitive function or other. Specifically, food-related tasks included contrasts using food-related stimuli (including visual food images and gustatory tasting of sweet solutions). Body-related tasks included contrasts using body images and body shape descriptive words. Emotion-related tasks included contrasts using stimuli or paradigms with emotional valence (including facial expressions, words/descriptions with emotional or social valence, and reward-related stimuli/paradigms). Cognitive-function-related tasks included the Wisconsin Card Sorting Test (WCST), go/no-go test, n-back task, and others. “Other experiments” were those not categorized into the above 4 groups. Readers may reference a detailed description of task type classification of each study in Table S2 in the Supplementary Materials.

As reported by Eickhoff et al (11), at least 17 experiments are required to control the contribution of one experiment in ALE meta-analysis. However, due to the limited number of included studies and multiple task types, it was not possible to meet this criterion in some subgroup and/or task-specific meta-analyses (Table 1 and Table 2). To attain instructive information about these questions, we conducted meta-analyses in all datasets with ≥ 5 experiments. As described below, results derived from meta-analyses with less than 17 experiments are only exploratory and should be interpreted cautiously.

### Activation likelihood estimation

The principles and procedures of ALE meta-analysis and its evolution have been described previously (10–12). Relevant to the present analysis, meta-analytic methods have recently been created to control for the effect of multiple neighboring foci within one experiment (10) and to minimize the false-positive rate by adopting a cluster-level family-wise error (FWE) rate to correct for multiple comparisons (11). Therefore, we utilized the most up-to-date and appropriate methods for our dataset, which account for multiple experiments from one study and for limiting the effect of multiple foci from one experiment biasing the overall meta-analysis. We used a cluster-level family-wise-error-rate-corrected (FWER-corrected) threshold of *P* < 0.05 with uncorrected voxel-level threshold of *P* < 0.001 for all meta-analyses. We performed all meta-analyses within MNI152 coordinate space. For a detailed description of the ALE procedures used in this study, please refer to the supplementary materials.

## Results

Our literature search and screening process follows the Preferred Reporting Items for Systematic Reviews and Meta-analyses (PRISMA, see Figure 1). A detailed description of our search strategy and studies excluded after a full-text assessment is shown in Supplementary Materials. Sixty-three studies (27 AN-C vs. HC, 11 AN-R vs. HC, 5 AN-C + AN-R, 8 AN+BN, 12 BN) with 101 experiments (76 AN vs. HC, 25 BN vs. HC), consisting of 1289 patients (703 AN-C, 297 AN-R and 289 BN) were included in the meta-analyses.

### Disorder-level meta-analyses: ED (AN+BN) vs. HC across all tasks

We conducted a ED vs. HC group meta-analysis of 101 experiments (76 AN, 25 BN) with 569 foci. We observed significant convergence of brain activity alteration in the right fusiform, right declive and left IPL in ED relative to HC groups (Table 1; Figure 2A). In contrast analyses, the ED group showed significant convergence of hypoactivation in the left precuneus (Table 1; Figure 2A). No cluster of ED-related hyperactivation passed our statistical threshold. (Given the large number of meta-analyses, ALE scores, ALE cluster volumes, and centers of mass are reserved for the Supplementary Materials.)

**Figure 2.**
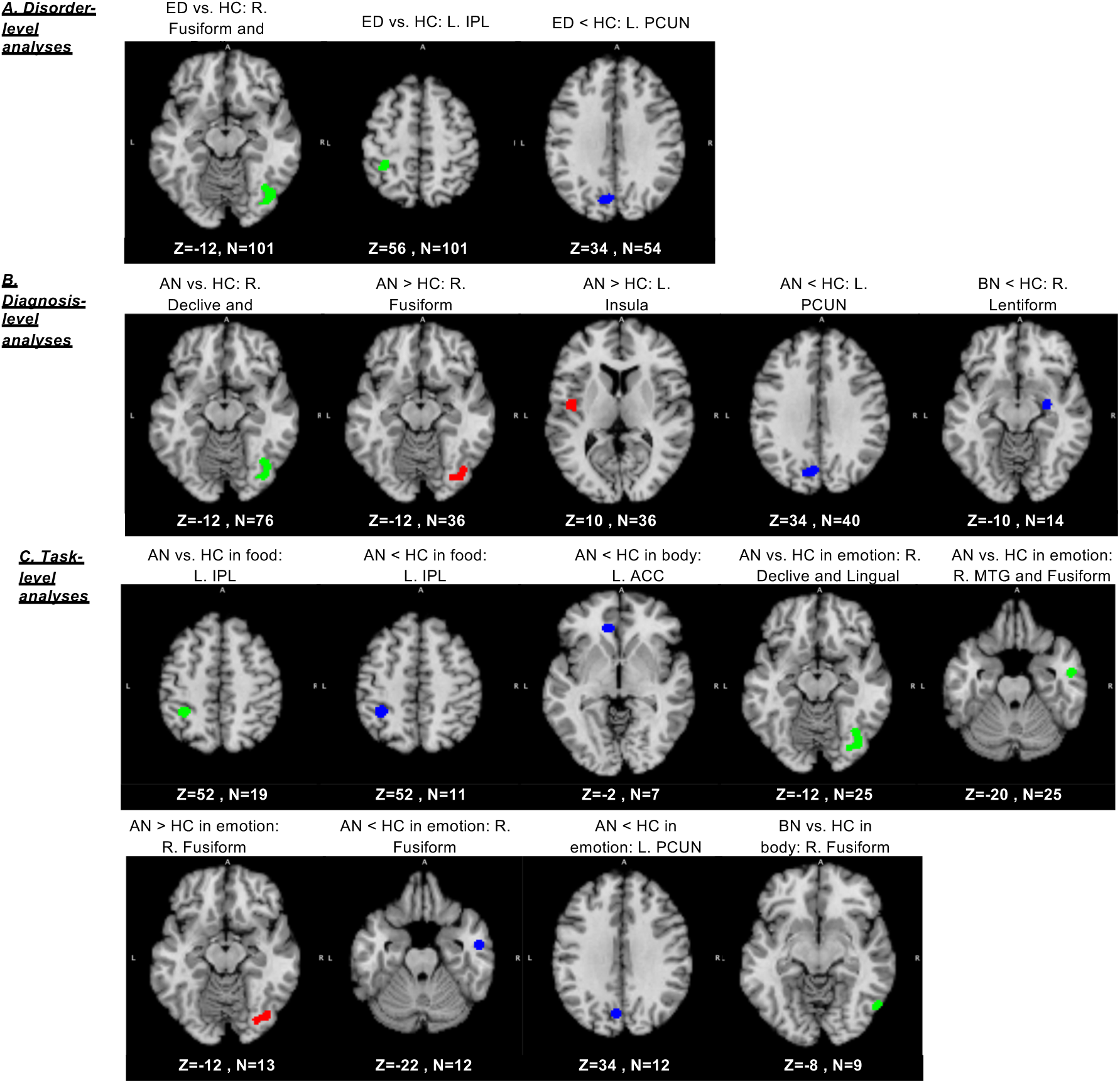
Altered brain function in AN and BN in hierarchical meta-analyses. Green, Red and Blue represents results from vs. HC, >HC and <HC analyses, respectively. Z indicates the Z coordinate of each axial slice. N indicates the number of experiments included in each meta-analysis. Abbreviations: ED: Eating Disorder; AN: Anorexia Nervosa; BN: Bulimia Nervosa; HC: Healthy Control; R: right; L: left; PCUN: Precuneus; ACC: Anterior Cingulate Cortex; IPL: Inferior Parietal Lobule; MTG: Middle Temporal Gyrus.

### Diagnosis-level meta-analyses: AN/BN vs. HC across all tasks

Results for AN vs. HC and AN < HC meta-analyses largely overlapped with the results of ED vs. HC and ED < HC meta-analyses (Table 1; Figure 2B). The AN group also showed significant convergence of hyperactivation in the right fusiform gyrus and left insula (Table 1; Figure 2B), which we did not observe in the ED>HC analysis. No significant cluster was observed in the BN vs. HC or BN > HC analyses, but the BN < HC analysis revealed a significant cluster in the right lentiform (right lateral globus pallidus nucleus) (Table 1; Figure 2B).

We were unable to replicate the above findings in any of the diagnosis-specific *sub-analyses* (i.e. AN-R, rAN, comorbidity-free or medication-free AN/BN; see Table 2 and Figure 3). Since most of the included studies used correction for multiple comparisons, the findings of sub-analyses for corrected studies largely mirrored the diagnosis-level meta-analyses for AN or BN (Table 2; Figure 3), with the exception of the BN < HC analysis, which revealed no significant cluster in this sub-analysis for corrected experiments (Table 2).

**Figure 3.**
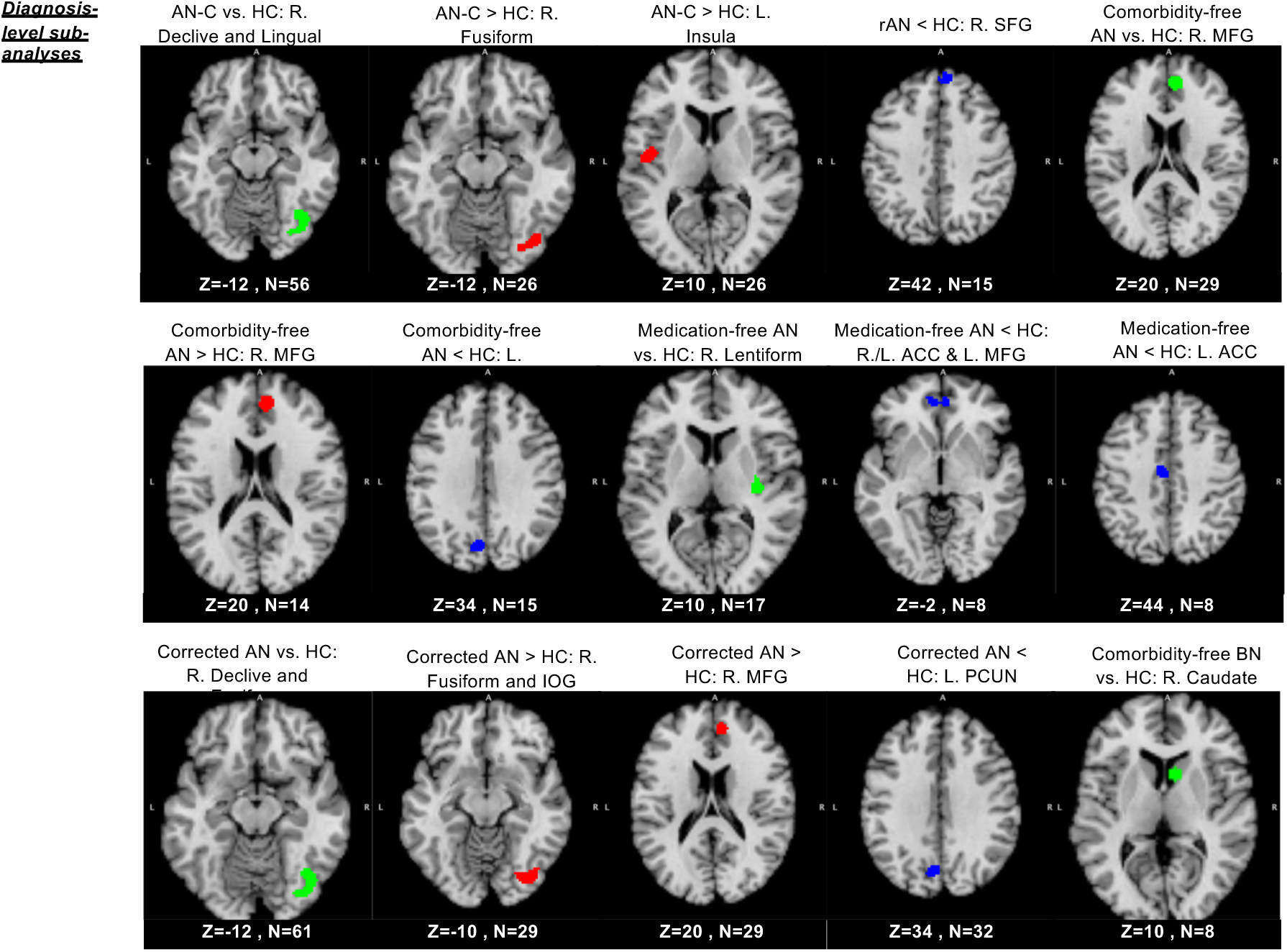
Altered brain function in AN and BN in diagnosis-level sub-analyses. Green, Red and Blue represents results from vs. HC, >HC and <HC analyses, respectively. Z indicates the Z coordinate of each axial slice. N indicates the number of experiments included in each meta-analysis. Abbreviations: AN: Anorexia Nervosa; BN: Bulimia Nervosa; HC: Healthy Control; rAN: restrictive AN; R: right; L: left; SFG: Superior Frontal Gyrus; MFG: Medial Frontal Gyrus; PCUN: Precuneus; ACC: Anterior Cingulate Cortex; IOG: Inferior Occipital Gyrus.

### Task-level meta-analyses: food, body, emotion and cognitive function

In food- and body-related tasks, the AN group showed consistent hypoactivation in the left IPL and in the left anterior cingulate cortex (ACC), respectively (see Table 1; Figure 2C). In emotion-related tasks, the AN vs. HC, AN>HC and AN<HC results largely mirrored those of the disease-level AN vs. HC, AN > HC and AN < HC meta-analyses (which were across all tasks), suggesting that the findings of diagnosis-level meta-analyses of AN were mainly related to the emotion-related tasks (Table 1; Figure 2). Cognitive-related tasks showed no statistically significant convergence in any of the AN groups. In BN studies, due to the limited number of included studies (<17), we conducted only 4 exploratory task-level meta-analyses: BN vs. HC and BN > HC in food-related tasks; BN vs. HC and BN < HC in body-related tasks. Only the BN vs. HC contrast in body-related tasks showed a statistically significant convergence, implicating the right fusiform gyrus (Table 1; Figure 2C).

## Discussion

The task-based functional MRI literature in AN and BN is quite large and reflects a diverse implementation of cognitive tasks across a clinically heterogenous patient group. To most ably synthesize this heterogeneity, we performed a hierarchical series of 49 total meta-analyses. Across all 101 experiments meta-analyzed at the disorder level (AN + BN), only the right fusiform gyrus, left IPL, and left precuneus survived rigorous statistical thresholds. These regions were not consistently observed across all hierarchies. Subsequent analyses at the diagnosis-level indicated that these overall results largely reflected differences observed in AN patients relative to HCs; specifically, the right fusiform gyrus was more activated in AN than HC and the left precuneus was less active in AN than HC. Specific task-level analyses recapitulated the findings implicating the right fusiform and left precuneus and the left IPL observed in other analytic hierarchies. These results are further understood in context of the diagnosis-level sub-meta-analyses, which indicated convergence limited to patients with current (as opposed to recovered) AN, again within the right fusiform gyrus. Given constraints of the current literature, some of our meta-analyses should be considered exploratory (those representing <17 experiments); however, overall our hierarchical analyses suggest that the most consistent results are found across studies of patients with current AN that employed either a food- or emotion-related task. Below, we discuss how this heterogeneity may reflect concerns present throughout functional brain imaging and what this substantial series of analyses means for future task-based fMRI studies of EDs.

### The neurophysiology of AN and BN

Previous reviews of the resting-state fMRI literature have suggested that dysconnectivity within corticolimbic circuits was related to impaired cognitive control and body image disturbances seen in EDs (13). Our results largely diverge from this model; notwithstanding the large task-based fMRI literature drawn upon in our analysis (101 experiments), our results of consistent cross-study convergence were limited to the right fusiform gyrus, left IPL, and left precuneus.

The fusiform gyrus comprises two sub-regions: the fusiform body area and the fusiform face area, which are implicated in processing of human body (14, 15) and facial expressions (16), respectively. The fusiform is thought to be a key node of the social information processing network, specifically in organizing information about one’s own and other’s facial expressions (17). The fusiform gyrus is also associated with preliminary perceptual processing of body/shape and facial expression/social stimuli (17). Previous studies have hypothesized that hyperactivation of the fusiform gyrus in AN may indicate elevated salience in detecting body/shape images and facial expressions, which is consistent with distorted body/shape perception (1) and increased sensitivity to punishment in social contexts (18); however our study did not find evidence supporting this claim, perhaps due to the small sample-size in the body-related tasks.

The IPL and the nearby temporo-parietal junction are associated with appetite regulation (19, 20), hunger state, and desire for food (21). Thus, dysfunction in the IPL and temporo-parietal junction may serve as another hub wherein dysfunction is associated with EDs.

The precuneus is associated with positive social information processing. Hypoactivation in the precuneus has been implicated in the decreased evaluation of positive social information in AN, which is consistent with the social anxiety and social avoidance symptoms of AN (22). A previous structural MRI study has shown reduced grey matter in the precuneus in AN (23); however, this finding could be confounded by malnourishment status (3).

We note that each of the functional mapping studies mentioned above associates a brain region with a cognitive task that has behavioral relevance (at least in theory) to EDs. While this suggests that such tasks might effectively engage brain circuits implicated in EDs, this should be viewed with caution, as discussed below.

### The complexity of clinical task-based fMRI studies

All experiments included in our meta-analysis were whole-brain task-based fMRI studies. Task-based functional MRI allows experimentally-induced perterburations of brain activity (measured indirectly with blood flow) within task-specific circuits to be related to experimental groups (in this case AN or BN). Our results are notable because there was no clear consensus across all meta-analytic hierarchies. Our results further indicate that the AN and BN literature does not report a common dysfunctioning circuit across food-related, emotion-related, and cognitive tasks. Importantly, we do not view our results as a failure of the task-based fMRI procedure in EDs; different tasks are designed to engage different brain circuits. We suspect these results likely reflect heterogeneity at the task and patient level.

Cognitive tasks are complex to design and implement. Variablity in the specific cognitive concept tested (e.g. type of emotional task), the precise task design (e.g. stimulus type, sequence, duration, and context), and the implementation of similar or identical tasks in separate laboratories (e.g. using different projectors or pulse sequences) represent potential confounding variables that may complicate across-study comparison. Indeed, from pre- to post-processing of fMRI data, the statistical analyses performed in individual studies can result in divergent and potentially false-positive results (24). Coordinate-based meta-analysis such as we have performed constitute arguably the most practical and statistically rigorous method to tease apart false-from true-positive findings across the published literature. Yet, even if task design and processing pipelines were perfectly standardized across laboratories, if clinical diagnosis remained the only group criterion (e.g. “AN” or “HC”), substantial cognitive heterogeneity would still likely exist within and across groups, and thus may yield inconsistent results across studies.

### EDs as a clinical continuum

Multiple converging lines of evidence suggest that EDs should not be viewed as a binary diagnostic group (i.e., AN or HC), but rather that patients should be placed on a continuum of nutritional and cognitive status. Structural MRI studies, for example, have strongly suggested that malnourishment (including food and water deprivation) status within AN relates strongly to differences in brain structure (3), something that cannot be captured by a diagnostic label. This is consistent with a previous meta-analysis of morphological MRI studies showing restoring of gray-matter and white-matter differences in AN after 1.5-8 years of clinical remission (25). In addition to the effect of nutritional status on morphological differences, a previous review of functional neuroimaging studies of AN also found that the brain functional alterations in AN are largely normalized after recovery (26).

Some authors have argued for a trait-based model wherein individuals could be placed on a spectrum of behavior (e.g. ‘impulse control’) that is more likely to represent ED’s underlying neuropathophysiology (27). Such an effort could abrogate the need for a diagnostically “pure” AN or BN (i.e., an ED in the absence of other psychiatric comorbidities, which is clinically (extremely) rare). A trait-based model would further place EDs more ably on a Research Domain Criterion (RDOC) framework (28). Until neurophysiologically meaningful experimental groups are included as part of the experiemental design, substantial cross-study convergence of neurophysiological results seems unlikely.

### In context with previous studies

Our results largely do not replicate a previous coordinate-based meta-analyses (9). Zhu et al (9) reported AN hyperactivation (i.e., AN>HCs) in emotion-related regions (frontal, caudate, uncus, insula and temporal) and hypoactivation in parietal cortex in processing of food- and body-related stimuli (Figure S1). The differences may relate to several factors. First, while our study included 76 whole-brain experiments of AN, Zhu et al (9) included only 21 experiments (9 food-, 8 body- and 4 emotion-related tasks). The limited number of experiments may have made it vulnerable to false-positive findings (11). Second, we applied the latest coordinate-based meta-analytic methods, including an improved cluster-level correction of multiple comparisons that is more sensitive to false-positive errors. Third, our hierarchical analysis suggests that disorder-level results were mainly driven by emotion-related tasks, which differs from the prior study (9) that was mainly limited to studies of food- and body-related stimuli. Notwithstanding these differences, our results replicate the AN-related hypoactivation in the left IPL in food-related tasks reported previously (9), even though the literature as a whole did not confirm other results. It is possible that hypoactivity of the left IPL (specifically in food-related tasks) may be a feature shared by individuals across the AN continuum and may serve as a central hub for brain dysfunction that leads to decreased appetite and overcontrol in food intake in AN.

### Limitations

Although this is the largest meta-analysis of AN to date and, to the best of our knowledge, the first meta-analysis of BN, it is prudent to enumerate the limitations of our analyses and why our results should be interpreted cautiously. First, many sub-analyses were limited by the relatively small number of experiments found in the literature. Any meta-analysis with <17 experiments should be interpreted cautiously, as such meta-analyses may be considered insufficiently powered. We present such results as exploratory and preliminary analyses. Second, due to the limitations of ALE meta-analyses, we are unable to conduct a publication bias and sensitivity analysis for the significant clusters. We further are unable to conduct regression analysis of potential confounding factors, like age, body mass index, duration of illness, sample size, and others. Third, given limitations in the published literature, we restricted our analyses to AN and BN at the exclusion of other eating disorders like binge-eating disorder or eating disorder not otherwise specified. Fourth, we grouped together in domains different tasks, and these may assess to varying degrees different aspects within the specific domains.

### Future directions

Our results draw attention to the influence that clinical population and task type can have on task-based fMRI analyses. When planning and comparing task-based fMRI studies in EDs, therefore, careful attention should be given to the clinical populations and tasks being utilized. Moreover, the interpretation of the findings in one single study should be performed cautiously and considered as representing a specific population under a specific task condition. Given the significant literature showing differences in brain structure and function associated with nutritional (and even prandial) status, it would be prudent for future studies to carefully characterize patients based on nutritional status. In addition, adopting a trait- or cognitive-domain-based approach to categorizing clinical patients with an ED (28) may more likely provide consistent neurophysiological findings than an approach focused on a constellation of symptoms.

In conclusion, the task-based functional MRI literature in AN and BN represents a heterogenous set of tasks and patients. Our hierarchical series of 49 total meta-analyses indicates that across 101 total experiments, the most consistent results are found within studies of patients with current AN that employed either a food- or emotion-related task. We recommend that future studies of EDs carefully characterize patients based on nutritional status and utilize a trait- or cognitive-domain-based approach to supplement clinical diagnosis.

## Disclosures and acknowledgments

This study was supported by the following agencies: China Scholarship Council (201706370081) to Bangshan Liu and (201706370272) to Yumeng Ju; the National Science and Technologic Program of China (2015BAI13B02) to Lingjiang Li; the National Basic Research Program of China (+ 2013CB835100) to Lingjiang Li; the National Natural Science Foundation of China (81171286 & 91232714) to Lingjiang Li and (81671353) to Yan Zhang; the U.S. National Institutes of Health (T32 MH019961, R25 MH071584) to Daniel S. Barron. We report no conflicts of interest with respect to the content of this manuscript.

We offer our sincere gratitude to Prof. Yufeng Zang from Hangzhou Normal University and Prof. Lin Xu from Chinese Academy of Sciences for their valuable comments and suggestions on the format and contents of this paper. We also thank Prof. Henry Yiyun Huang from Yale University for his administrative assistance in providing studying space and organizing member collaboration for our work.

