## Supplementary Materials for "Altered brain function in anorexia nervosa and bulimia nervosa: A hierarchical series of task-based fMRI meta-analyses"

**Literature search**

**(1) Electronic databases searching**

1) Pubmed (2018/01/21)

(((((neuroimaging[Title/Abstract] or magnetic resonance imaging[Title/Abstract] or MRI[Title/Abstract] or fmri[Title/Abstract] OR functional magnetic imaging[Title/Abstract]))) AND (((anorexia nervosa[Title] OR bulimia nervosa[Title] OR eating disorder[Title] OR eating disorders[Title]))))

Results: 359 items;

2) Web of Science (2018/01/21)

TOPIC: (neuroimaging or magnetic resonance imaging or MRI or fmri OR functional magnetic imaging) AND TITLE: (anorexia nervosa OR bulimia nervosa OR binge eating disorder OR eating disorder OR eating disorders)

Timespan: All years. Indexes: SCI-EXPANDED, SSCI, A&HCI, CPCI-S, CPCI-SSH, BKCI-S, BKCI-SSH, ESCI, CCR-EXPANDED, IC.

Results: 439 items

3) Embase (2018/01/22)

Title or Abstract: (neuroimaging or magnetic resonance imaging or MRI or fmri OR functional magnetic imaging) AND Title: (anorexia nervosa OR bulimia nervosa OR eating disorder OR eating disorders)

Results: 506 items

4) PsycINFO (2018/01/23) (EBSCO-PsycINFO)

AB ( neuroimaging or magnetic resonance imaging or MRI or fmri OR functional magnetic imaging ) AND TI ( anorexia nervosa OR bulimia nervosa OR eating disorder OR eating disorders ) Source Type: Academic Journals or Dissertations.

Results: 276 items

5) Chinese databases (including China National Knowledge Infrastructure (CNKI), Chinese Biomedical Literature Database (CBM), and WANFANG database): 2018/01/22

Search terms: AB ( neuroimaging or magnetic resonance imaging or MRI or fmri OR functional magnetic imaging ) AND TI ( anorexia nervosa OR bulimia nervosa OR anorexia OR bulimia OR eating disorder OR eating disorders ) . In Chinese: “神经性厌食”or“神经性贪食”or“厌食症”or“贪食症”or“暴食症”or“进食障碍”AND “功能磁共振” or “神经影像”

Results:

CNKI: 5 items; CBM: 20 items; WANFANG: 4 items. In total: 5 + 20 + 4 = 29 items

References retrieved from online database searching: 359+439+506+276+29 = 1609 items

**(2) Manual searching**

We searched the references of 5 relevant published reviews or meta-analyses(1-5). The manual searching attached 72 items.

**Literatures screening**

The process of literature screening was shown in Fig 1 in the manuscript. Fifty papers were excluded in the full-text screening for the following reasons:

1) Studies with other neuroimaging techniques rather than fMRI (6-8) or studies using resting-state fMRI (9) or studies using multiple modal neuroimaging technique and no independent fMRI data was available (10)

2) Studies using Binge eating disorder subjects or other eating disorders (11, 12)

3) Studies with no healthy comparison group (13-15) or sample size too small (n<5 in any group) (16)

4) Studies with no direct between-group comparison (17-30)

5) Studies with only region of interest (ROI) or seed-based analyses or using small volume correction (SVC) for multiple comparisons (31-42)

6) Studies showing no significant group difference (43-52)

7) No coordinates were reported in the article, and the coordinates cannot be acquired through contacting with the authors (53-55)

**Records of author-contacting during the literature screening and data extraction process**

**Table S1. Summary of author-contacting information in the data collection process**

| Study | Contacted authors & emails | Reason | Result |
| --- | --- | --- | --- |
| (53) | V. Ricca  | Ask for the unreported coordinates of the “gravity center” | Responded. The authors could no longer find the data. Thus, this study was finally excluded. |
| (56) | Kate Tchanturia | Ask for information about the gender distribution of the sample | Responded. All participants were female. |
| (57) | W. H. Kaye   Psychological Medicine customer service  | Ask for the online supplement where the coordinates were reported | Responded by the journal customer service staff. The coordinates were extracted from the online supplement. This paper was finally included. |
| (55) | Yuan Shen  | Ask for the unreported coordinates of the cluster showing significant difference between BN and HC. | No response. The paper was finally excluded due to lack of data for analysis. |

We also contacted the authors of conference abstracts to inquire whether their data has been published in journal articles or whether they could share their data with us, the conference abstracts include: (58-66). This process helped us attach a new journal article(67), which was included in our meta-analysis.

**Activation likelihood estimation (ALE) meta-analyses**

Activation likelihood estimation (ALE) is a coordinate-based meta-analytic method that quantifies the convergence of task-based fMRI experiments. The ALE algorithm has evolved over time to address specific limitations and to meet the computational demands of the community (68). An “experiment” in ALE meta-analysis is defined as the contrast that involves the same or similar brain function in one study. Thus, one published study may have several “experiments” since a task may involve different brain function. There are three major steps in ALE meta-analysis. First, the coordinates extracted from the included studies are converted to three-dimensional gaussian distributions with a full width half maximum determined by the smallest group sample size of the experiment. Our meta-analyses were conducted in MNI space, with the coordinates originally reported in Talairach space converted by the Lancaster transformation. MNI coordinates attained from the Brett transformation were converted back to Talaraich space and then reconverted to MNI space with the Lancaster transformation. Second, a whole-brain map of voxel-wise probability of activation is generated with each voxel assigned an ALE value in each experiment. The ALE value in each voxel is determined by the gaussian kernel with the highest probability of activation in each experiment. Third, the ALE maps of the experiments are merged to detect the convergence of probability of activation. Specifically, the ALE values in different experiments are compared with a null distribution created by permutation analysis to identify randomized clusters (noise) and nonrandomized clusters (signal). A cluster-level family-wise error rate (FWER) corrected threshold of *P* < 0.05 with uncorrected voxel-level threshold of *P* < 0.001 was selected as the threshold for all meta-analyses in this study. All the ALE meta-analyses were conducted in GingerALE 2.3.6 for MacOS system. The results were read with Mango 4.0.1 for MacOS system with images overlaid on a Collins27 T1 MNI template.

**Basic information about the included studies**

The characteristics of the included studies are shown in Table S2 (attached below).

**Comprehensive results of hierarchical meta-analyses and diagnosis-level sub-analyses**

**Table S3. Statistical Results for Hierarchical Meta-Analyses.**

|  |  |  |  |  | weighted Center^*^ | | |  | maximum ALE value^*^ | | |  |
| --- | --- | --- | --- | --- | --- | --- | --- | --- | --- | --- | --- | --- |
| Analysis and Cluster number | | # experiments | # foci | Volume (mm^3^) | x | y | z | Extrema Value (X 10^2) | x | y | z | Anatomical label |
| **Disorder-level analysis** | | | | | |  |  |  |  |  |  |  |
| ED vs. HC | | 101 | 569 |  |  |  |  |  |  |  |  |  |
| 1 | |  |  | 1616 | 39.5 | -69.9 | -12.8 | 2.69 | 42 | -70 | -14 | Right Cerebrum.Occipital Lobe.Fusiform Gyrus.Gray Matter.Brodmann area 19 |
|  | |  |  |  |  |  |  | 2.39 | 38 | -64 | -14 | Right Cerebellum.Posterior Lobe.Declive.Gray Matter.* |
|  | |  |  |  |  |  |  | 2.12 | 36 | -78 | -8 | Right Cerebrum.Occipital Lobe.Fusiform Gyrus.Gray Matter.Brodmann area 19 |
| 2 | |  |  | 808 | -35.5 | -41.8 | 53.2 | 2.28 | -34 | -42 | 52 | Left Cerebrum.Parietal Lobe.Inferior Parietal Lobule.Gray Matter.Brodmann area 40 |
| ED>HC | | 47 | 290 |  |  |  |  |  |  |  |  |  |
| None | |  |  |  |  |  |  |  |  |  |  |  |
| ED<HC | | 54 | 279 |  |  |  |  |  |  |  |  |  |
| 1 | |  |  | 752 | -7.9 | -72.5 | 36.7 | 1.87 | -6 | -72 | 34 | Left Cerebrum.Occipital Lobe.Precuneus.Gray Matter.Brodmann area 31 |
|  | |  |  |  |  |  |  | 1.73 | -6 | -72 | 42 | Left Cerebrum.Parietal Lobe.Precuneus.Gray Matter.Brodmann area 7 |
| **Diagnosis-level analyses** | | | | |  |  |  |  |  |  |  |  |
| AN vs. HC | | 76 | 458 |  |  |  |  |  |  |  |  |  |
| 1 | |  |  | 1592 | 38.2 | -70.3 | -12.4 | 2.33 | 38 | -64 | -14 | Right Cerebellum.Posterior Lobe.Declive.Gray Matter.* |
|  | |  |  |  |  |  |  | 2.11 | 36 | -78 | -8 | Right Cerebrum.Occipital Lobe.Fusiform Gyrus.Gray Matter.Brodmann area 19 |
|  | |  |  |  |  |  |  | 2.08 | 42 | -68 | -14 | Right Cerebellum.Posterior Lobe.Declive.Gray Matter.* |
| AN>HC | | 36 | 251 |  |  |  |  |  |  |  |  |  |
| 1 | |  |  | 840 | 36.8 | -75.6 | -11.1 | 1.66 | 38 | -76 | -12 | Right Cerebrum.Occipital Lobe.Fusiform Gyrus.Gray Matter.Brodmann area 19 |
| 2 | |  |  | 728 | -45.7 | -8.8 | 10.1 | 1.79 | -46 | -8 | 10 | Left Cerebrum.Sub-lobar.Insula.Gray Matter.Brodmann area 13 |
| AN<HC | | 40 | 207 |  |  |  |  |  |  |  |  |  |
| 1 | |  |  | 984 | -8.1 | -72.5 | 36.4 | 1.87 | -6 | -72 | 34 | Left Cerebrum.Occipital Lobe.Precuneus.Gray Matter.Brodmann area 31 |
|  | |  |  |  |  |  |  | 1.72 | -6 | -72 | 42 | Left Cerebrum.Parietal Lobe.Precuneus.Gray Matter.Brodmann area 7 |
| BN vs. HC | | 25 | 111 |  |  |  |  |  |  |  |  |  |
| None | |  |  |  |  |  |  |  |  |  |  |  |
| BN>HC | | 11 | 39 |  |  |  |  |  |  |  |  |  |
| None | |  |  |  |  |  |  |  |  |  |  |  |
| BN<HC | | 14 | 72 |  |  |  |  |  |  |  |  |  |
| 1 | |  |  | 992 | 27.3 | -7.9 | -8.3 | 1.40 | 28 | -8 | -10 | Right Cerebrum.Sub-lobar.Lentiform Nucleus.Gray Matter.Lateral Globus Pallidus |
|  | |  |  |  |  |  |  | 0.90 | 22 | -6 | 0 | Right Cerebrum.Sub-lobar.Lentiform Nucleus.Gray Matter.Lateral Globus Pallidus |
| **Task-level analyses** | | | | |  |  |  |  |  |  |  |  |
| AN vs. HC food | | 19 | 71 |  |  |  |  |  |  |  |  |  |
| 1 | |  |  | 792 | -35.7 | -43.3 | 52 | 1.38 | -36 | -44 | 52 | Left Cerebrum.Parietal Lobe.Inferior Parietal Lobule.Gray Matter.Brodmann area 40 |
| AN>HC food | | 8 | 36 |  |  |  |  |  |  |  |  |  |
| None | |  |  |  |  |  |  |  |  |  |  |  |
| AN<HC food | | 11 | 35 |  |  |  |  |  |  |  |  |  |
| 1 | |  |  | 1008 | -35.8 | -43.3 | 52 | 1.38 | -36 | -44 | 52 | Left Cerebrum.Parietal Lobe.Inferior Parietal Lobule.Gray Matter.Brodmann area 40 |
| AN vs. HC body | | 12 | 100 |  |  |  |  |  |  |  |  |  |
| None | |  |  |  |  |  |  |  |  |  |  |  |
| AN>HC body | | 5 | 69 |  |  |  |  |  |  |  |  |  |
| None | |  |  |  |  |  |  |  |  |  |  |  |
| AN<HC body | | 7 | 31 |  |  |  |  |  |  |  |  |  |
| 1 | |  |  | 568 | -10 | 37 | -3.8 | 0.94 | -8 | 36 | -4 | Left Cerebrum.Limbic Lobe.Anterior Cingulate.Gray Matter.Brodmann area 24 |
| AN vs. HC emotion | | 25 | 135 |  |  |  |  |  |  |  |  |  |
| 1 | |  |  | 1880 | 36.3 | -72 | -10.9 | 1.94 | 36 | -64 | -14 | Right Cerebellum.Posterior Lobe.Declive.Gray Matter.* |
|  | |  |  |  |  |  |  | 1.58 | 36 | -76 | -8 | Right Cerebrum.Occipital Lobe.Lingual Gyrus.Gray Matter.Brodmann area 18 |
| 2 | |  |  | 832 | 54.2 | -5.6 | -23.4 | 1.85 | 54 | -6 | -22 | Right Cerebrum.Temporal Lobe.Middle Temporal Gyrus.Gray Matter.Brodmann area 21 |
|  | |  |  |  |  |  |  | 1.65 | 56 | -6 | -30 | Right Cerebrum.Temporal Lobe.Fusiform Gyrus.Gray Matter.Brodmann area 20 |
| AN>HC emotion | | 13 | 61 |  |  |  |  |  |  |  |  |  |
| 1 | |  |  | 720 | 34.3 | -76.2 | -11.1 | 1.24 | 30 | -78 | -10 | Right Cerebrum.Occipital Lobe.Fusiform Gyrus.Gray Matter.Brodmann area 19 |
|  | |  |  |  |  |  |  | 1.17 | 38 | -76 | -12 | Right Cerebrum.Occipital Lobe.Fusiform Gyrus.Gray Matter.Brodmann area 19 |
| AN<HC emotion | | 12 | 74 |  |  |  |  |  |  |  |  |  |
| 1 | |  |  | 896 | 55 | -6 | -26 | 1.65 | 55 | -6 | -26 | Right Cerebrum.Temporal Lobe.Fusiform Gyrus.Gray Matter.Brodmann area 20 |
| 2 | |  |  | 872 | -6 | -72 | 38 | 1.67 | -6 | -72 | 38 | Left Cerebrum.Occipital Lobe.Precuneus.Gray Matter.Brodmann area 31 |
| AN vs. HC cognitive function | | 13 | 67 |  |  |  |  |  |  |  |  |  |
| None | |  |  |  |  |  |  |  |  |  |  |  |
| AN>HC cognitive function | | 6 | 19 |  |  |  |  |  |  |  |  |  |
| None | |  |  |  |  |  |  |  |  |  |  |  |
| AN<HC cognitive function | | 7 | 48 |  |  |  |  |  |  |  |  |  |
| None | |  |  |  |  |  |  |  |  |  |  |  |
| BN vs. HC food | | 9 | 30 |  |  |  |  |  |  |  |  |  |
| None | |  |  |  |  |  |  |  |  |  |  |  |
| BN>HC food | | 5 | 19 |  |  |  |  |  |  |  |  |  |
| None | |  |  |  |  |  |  |  |  |  |  |  |
| BN vs. HC body | | 9 | 30 |  |  |  |  |  |  |  |  |  |
| 1 | |  |  | 456 | 53.6 | -64.5 | -9.4 | 1.08 | 52 | -66 | -10 | Right Cerebrum.Temporal Lobe.Fusiform Gyrus.Gray Matter.Brodmann area 37 |
| BN<HC body | | 5 | 24 |  |  |  |  |  |  |  |  |  |
| None | |  |  |  |  |  |  |  |  |  |  |  |

Abbreviations: ED: Eating Disorder; AN: Anorexia Nervosa; BN: Bulimia Nervosa; HC: Healthy Control; None: no significant convergence.

**Table S4. Statistical Results for Diagnosis-Level Sub-Meta-Analyses.**

|  |  |  |  |  | weighted Center^*^ | | |  | maximum ALE value^*^ | | |  |
| --- | --- | --- | --- | --- | --- | --- | --- | --- | --- | --- | --- | --- |
| Analysis and Cluster number | | # experiments | # foci | Volume (mm^3^) | x | y | z | Extrema Value | x | y | z | Anatomical label |
| **Diagnosis-level sub-analyses** | | | | |  |  |  |  |  |  |  |  |
| AN-C vs. HC | | 56 | 285 |  |  |  |  |  |  |  |  |  |
| 1 | |  |  | 2000 | 38.6 | -68.8 | -13 | 2.33 | 38 | -64 | -14 | Right Cerebellum.Posterior Lobe.Declive.Gray Matter.* |
|  | |  |  |  |  |  |  | 2.07 | 42 | -68 | -14 | Right Cerebellum.Posterior Lobe.Declive.Gray Matter.* |
|  | |  |  |  |  |  |  | 1.60 | 36 | -76 | -8 | Right Cerebrum.Occipital Lobe.Lingual Gyrus.Gray Matter.Brodmann area 18 |
| AN-C>HC | | 26 | 140 |  |  |  |  |  |  |  |  |  |
| 1 | |  |  | 848 | 37 | -74.3 | -11.7 | 1.39 | 42 | -70 | -14 | Right Cerebrum.Occipital Lobe.Fusiform Gyrus.Gray Matter.Brodmann area 19 |
|  | |  |  |  |  |  |  | 1.36 | 40 | -74 | -12 | Right Cerebrum.Occipital Lobe.Fusiform Gyrus.Gray Matter.Brodmann area 19 |
|  | |  |  |  |  |  |  | 1.25 | 30 | -78 | -10 | Right Cerebrum.Occipital Lobe.Fusiform Gyrus.Gray Matter.Brodmann area 19 |
| 2 | |  |  | 776 | -46.6 | -7.6 | 9.5 | 1.70 | -46 | -6 | 10 | Left Cerebrum.Sub-lobar.Insula.Gray Matter.Brodmann area 13 |
| AN-C<HC | | 30 | 145 |  |  |  |  |  |  |  |  |  |
| None | |  |  |  |  |  |  |  |  |  |  |  |
| AN-R vs. HC | | 20 | 173 |  |  |  |  |  |  |  |  |  |
| None | |  |  |  |  |  |  |  |  |  |  |  |
| AN-R>HC | | 10 | 111 |  |  |  |  |  |  |  |  |  |
| None | |  |  |  |  |  |  |  |  |  |  |  |
| AN-R<HC | | 10 | 62 |  |  |  |  |  |  |  |  |  |
| None | |  |  |  |  |  |  |  |  |  |  |  |
| **Confounder analyses** | | | | |  |  |  |  |  |  |  |  |
| AN vs. HC all rAN | | 31 | 103 | 680 |  |  |  |  |  |  |  |  |
| None | |  |  |  |  |  |  |  |  |  |  |  |
| AN>HC all rAN | | 16 | 51 |  |  |  |  |  |  |  |  |  |
| None | |  |  |  |  |  |  |  |  |  |  |  |
| AN<HC all rAN | | 15 | 52 |  |  |  |  |  |  |  |  |  |
| 1 | |  |  | 800 | 6.3 | 51.1 | 40.3 | 1.21 | 6 | 52 | 42 | Right Cerebrum.Frontal Lobe.Superior Frontal Gyrus.Gray Matter.Brodmann area 8 |
| AN vs. HC Comorbidity free | | 29 | 198 | 640 |  |  |  |  |  |  |  |  |
| 1 | |  |  | 1104 | 7.2 | 47.7 | 20.4 | 2.69 | 8 | 48 | 20 | Right Cerebrum.Frontal Lobe.Medial Frontal Gyrus.Gray Matter.Brodmann area 9 |
| AN>HC Comorbidity-free | | 14 | 112 |  |  |  |  | . |  |  |  |  |
| 1 | |  |  | 992 | 6.9 | 47.6 | 20.3 | 1.88 | 8 | 48 | 20 | Right Cerebrum.Frontal Lobe.Medial Frontal Gyrus.Gray Matter.Brodmann area 9 |
| AN<HC Comorbidity-free | | 15 | 86 |  |  |  |  |  |  |  |  |  |
| 1 | |  |  | 1344 | -6.5 | -72.2 | 37.7 | 1.77 | -6 | -72 | 34 | Left Cerebrum.Occipital Lobe.Precuneus.Gray Matter.Brodmann area 31 |
|  | |  |  |  |  |  |  | 1.71 | -6 | -72 | 42 | Left Cerebrum.Parietal Lobe.Precuneus.Gray Matter.Brodmann area 7 |
| AN vs. HC Medication free | | 17 | 165 |  |  |  |  |  |  |  |  |  |
| 1 | |  |  | 728 | 33.4 | -16.1 | 10.3 | 1.55 | 34 | -16 | 10 | Right Cerebrum.Sub-lobar.Lentiform Nucleus.Gray Matter.Putamen |
| AN>HC Medication-free | | 9 | 105 |  |  |  |  |  |  |  |  |  |
| None | |  |  |  |  |  |  |  |  |  |  |  |
| AN<HC Medication-free | | 8 | 60 |  |  |  |  |  |  |  |  |  |
| 1 | |  |  | 816 | -2.1 | 49.7 | 0 | 1.08 | 4 | 50 | 0 | Right Cerebrum.Limbic Lobe.Anterior Cingulate.Gray Matter.Brodmann area 32 |
|  | |  |  |  |  |  |  | 0.90 | -12 | 52 | 2 | Left Cerebrum.Frontal Lobe.Medial Frontal Gyrus.Gray Matter.Brodmann area 10 |
|  | |  |  |  |  |  |  | 0.88 | -6 | 48 | -2 | Left Cerebrum.Limbic Lobe.Anterior Cingulate.Gray Matter.Brodmann area 32 |
| 2 | |  |  | 544 | -5.8 | -10.7 | 44.2 | 1.02 | -6 | -10 | 44 | Left Cerebrum.Limbic Lobe.Cingulate Gyrus.Gray Matter.Brodmann area 24 |
| AN vs. HC corrected | | 61 | 349 | 1104 |  |  |  |  |  |  |  |  |
| 1 | |  |  | 2024 | 38.2 | -70.6 | -12.3 | 2.33 | 38 | -64 | -14 | Right Cerebellum.Posterior Lobe.Declive.Gray Matter.* |
|  | |  |  |  |  |  |  | 2.11 | 36 | -78 | -8 | Right Cerebrum.Occipital Lobe.Fusiform Gyrus.Gray Matter.Brodmann area 19 |
|  | |  |  |  |  |  |  | 2.08 | 42 | -68 | -14 | Right Cerebellum.Posterior Lobe.Declive.Gray Matter.* |
| AN>HC corrected | | 29 | 178 |  |  |  |  |  |  |  |  |  |
| 1 | |  |  | 1296 | 36 | -75.8 | -10.8 | 1.66 | 38 | -76 | -12 | Right Cerebrum.Occipital Lobe.Fusiform Gyrus.Gray Matter.Brodmann area 19 |
|  | |  |  |  |  |  |  | 1.04 | 36 | -84 | -2 | Right Cerebrum.Occipital Lobe.Inferior Occipital Gyrus.Gray Matter.Brodmann area 18 |
| 2 | |  |  | 672 | 7 | 47.5 | 20.6 | 1.88 | 8 | 48 | 20 | Right Cerebrum.Frontal Lobe.Medial Frontal Gyrus.Gray Matter.Brodmann area 9 |
| AN<HC corrected | | 32 | 171 |  |  |  |  |  |  |  |  |  |
| 1 | |  |  | 848 | -6.4 | -72.2 | 36.9 | 1.81 | -6 | -72 | 34 | Left Cerebrum.Occipital Lobe.Precuneus.Gray Matter.Brodmann area 31 |
|  | |  |  |  |  |  |  | 1.71 | -6 | -72 | 42 | Left Cerebrum.Parietal Lobe.Precuneus.Gray Matter.Brodmann area 7 |
| BN vs. HC Comorbidity free | | 8 | 32 |  |  |  |  |  |  |  |  |  |
| 1 | |  |  | 640 | 8 | 13 | 9 | 1.42 | 8 | 12 | 10 | Right Cerebrum.Sub-lobar.Caudate.Gray Matter.Caudate Body |
| BN vs. HC Medication free | | 6 | 19 |  |  |  |  |  |  |  |  |  |
| None | |  |  |  |  |  |  |  |  |  |  |  |
| BN vs. HC corrected | | 19 | 84 |  |  |  |  |  |  |  |  |  |
| None | |  |  |  |  |  |  |  |  |  |  |  |
| BN>HC corrected | | 8 | 29 |  |  |  |  |  |  |  |  |  |
| None | |  |  |  |  |  |  |  |  |  |  |  |
| BN<HC corrected | | 11 | 55 |  |  |  |  |  |  |  |  |  |
| None | |  |  |  |  |  |  |  |  |  |  |  |

Abbreviations: AN-C: current Anorexia Nervosa; AN-R: recovered AN; rAN: restrictive AN; BN: Bulimia Nervosa; HC: Healthy Control; None: no significant convergence.

**Comparison between the results of our study and those of Zhu et al (2012)**

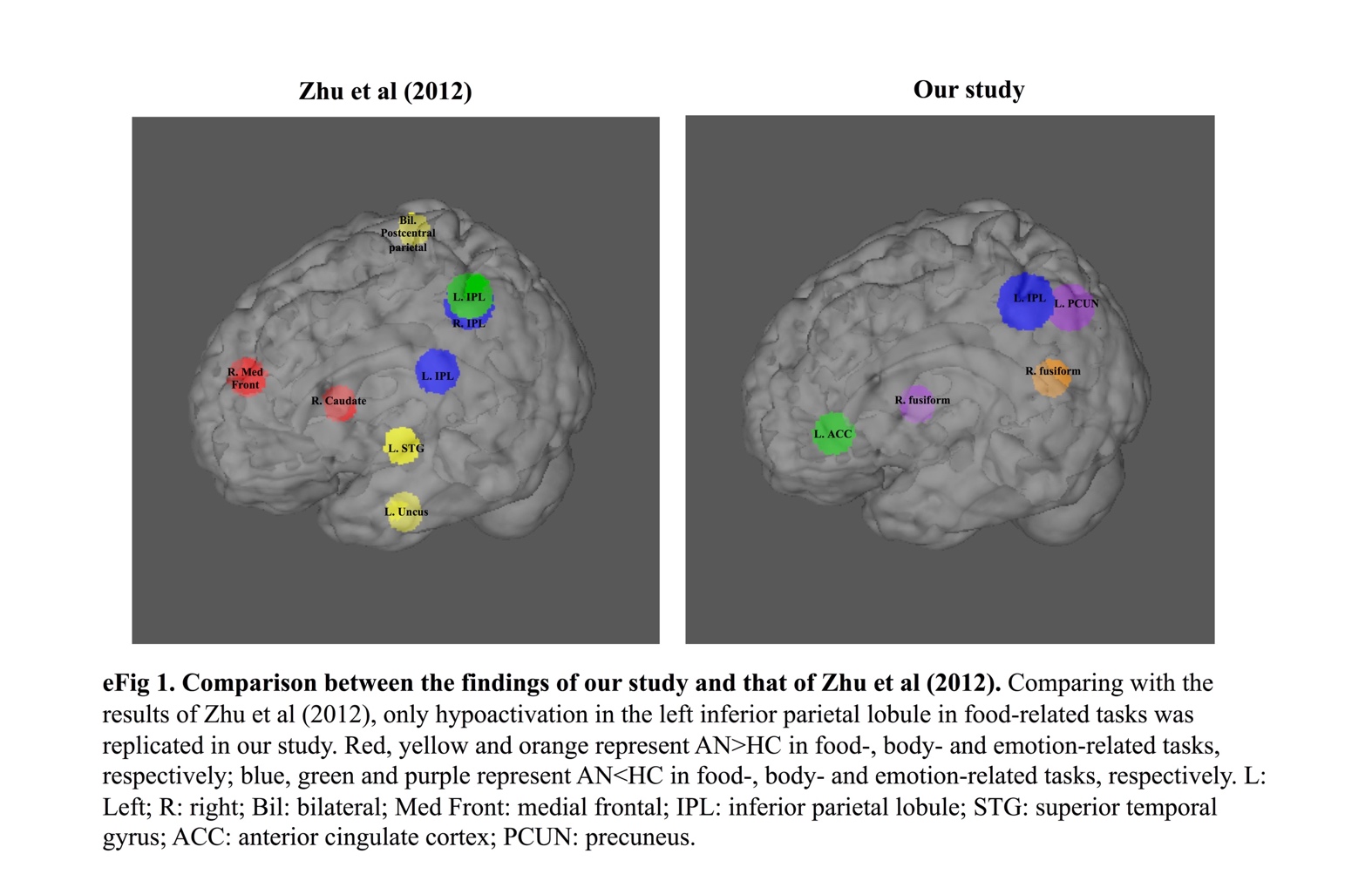

**Figure S1. Comparison between the results of our study and those of Zhu et al (2012).** Comparing with the results of Zhu et al (2012), only hypoactivation in the left inferior parietal lobule in food-related tasks was replicated in our study. Red, yellow and orange represent AN>HC in food-, body- and emotion-related tasks, respectively; blue, green and purple represent AN<HC in food-, body- and emotion-related tasks, respectively. L: Left; R: right; Bil: bilateral; Med Front: medial frontal; IPL: inferior parietal lobule; STG: superior temporal gyrus; ACC: anterior cingulate cortex; PCUN: precuneus.
